# Non-additive polygenic models improve predictions of fitness traits in three eukaryote model species

**DOI:** 10.1101/2020.07.14.194407

**Authors:** Moises Exposito-Alonso, Peter Wilton, Rasmus Nielsen

## Abstract

To describe a living organism it is often said that “the whole is greater than the sum of its parts”. In genetics, we may also think that the effect of multiple mutations on an organism is greater than their additive individual effect, a phenomenon called epistasis or multiplicity. Despite the last decade’s discovery that many disease- and fitness-related traits are polygenic, or controlled by many genetic variants, it is still debated whether the effects of individual genes combine additively or not. Here we develop a flexible likelihood framework for genome-wide associations to fit complex traits such as fitness under both additive and non-additive polygenic architectures. Analyses of simulated datasets under different true additive, multiplicative, or other epistatic models, confirm that our method can identify global non-additive selection. Applying the model to experimental datasets of wild type lines of *Arabidopsis thaliana, Drosophila melanogaster*, and *Saccharomyces cerevisiae*, we find that fitness is often best explained with non-additive polygenic models. Instead, a multiplicative polygenic model appears to better explain fitness in some experimental environments. The statistical models presented here have the potential to improve prediction of phenotypes, such as disease susceptibility, over the standard methods for calculating polygenic scores which assume additivity.

---

Over a decade of Genome-Wide Association (GWA) studies has confirmed what quantitative geneticists have long suspected - that many complex and continuous traits, including human health traits (1) and experimentally measured fitness in plants (2) and animals (3), are controlled by hundreds if not thousands of mutations (1). These observations suggest that adaptation might generally occur in a polygenic fashion, where multiple alleles in the genome are under selection, rather than through single selective sweeps, where a single allele contributes most of the heritability and has a large fitness effect (4). There are many possible ways in which multiple mutations can combine or interact to determine a trait. Still, the vast majority of polygenic models used for genetic mapping of traits exclusively consider additive effects, in which mutations are assumed to act independently on a trait, without any interaction. Considering the complex nature of biological systems (5), this is a counterintuitive assumption, which has generated controversy in quantitative and population genetics (6–8).

Quantitative genetics, the branch of evolutionary genetics focused on understanding the genetic contribution of traits, has GWA as one of its core tools. When conducting GWA using fitness or a fitness proxy as the trait of interest, one implicitly assumes selection is additive (i.e. fitness *w* = 1 + *s*_1_ + *s*_2_ + *s*_3_ …, with *s* being the selection coefficients or relative fitness effects of a given variant). The justification for this assumption is mostly statistical, and rooted in the origin of the field of quantitative genetics and the infinitesimal model (9), which posits that when many loci affect a trait in small quantities, even if there are interactions, the additive model is a good approximation — a theoretical notion which has been backed up by practical successes in artificial selection for breeding plant and animals (10–13).

Despite this success of the additive model, much classic and modern genetic research supports the existence of epistasis (14), encouraging researchers to explore this biological phenomenon in genome-wide or polygenic approaches. Some models in quantitative genetics have extended a linear model to include pairwise epistatic interactions include pairwise epistatic interactions (*w* = 1 + *s*_1_ + *s*_2_ + *s*_12_) (15, 16). Extending the pairwise epistasis model to genome-wide approaches, for a dataset *m* of SNPs, one would have to infer *m* main effects and *m*^2^ interaction terms, which likely is many more parameters than current datasets have the power to detect (16). Furthermore, admitti ng pairwise epistasis immediately leads to the question of higher-order epistasis, such as trios or quartets of interacting loci (17).

The pairwise epistatic model is only one of many non-additive models proposed in the broader field of evolutionary genetics (reviewed and discussed in (18) and (19)). Other important and commonly assumed models are: The “multiplicative model” (*w* = (1 + *s*_1_) (1 + *s*_2_)), where fitness increases in a geometric fashion with the number of alleles affecting fitness (a pairwise version of which also exist (20)). The “synergistic or positive epistasis model”, where fitness increases or decreases faster than a geometric function (21), and the “diminishing returns or negative epistasis model”, where fitness gains are less than expected from the combination of adaptive mutations and ultimately plateauing (**Fig. 1C**) (21, 22).

**Figure 1.**
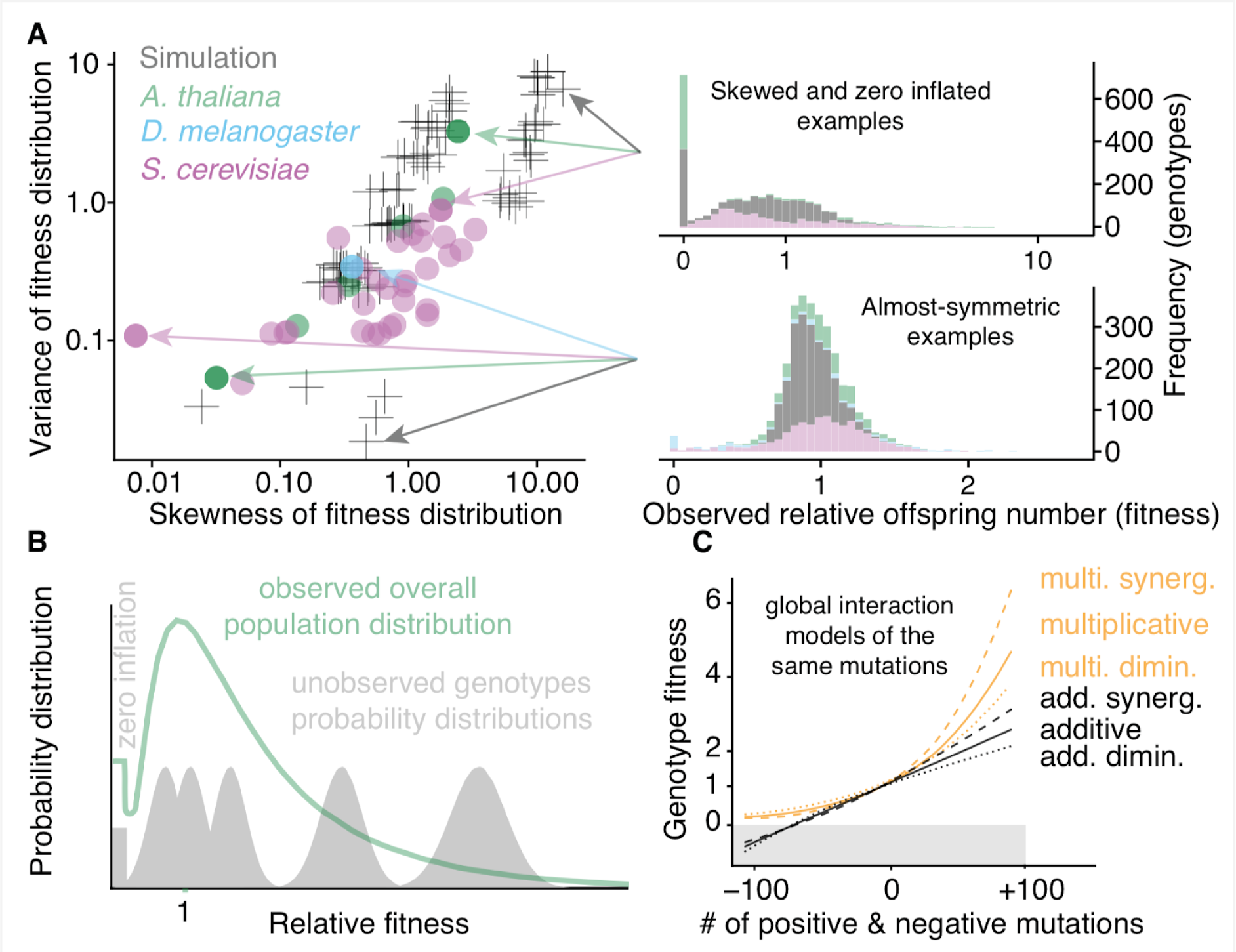
Modeling non-additive polygenic fitness. (**A**) Variance and skewness of relative offspring number (fitness) distributions of three species: Arabidopsis thaliana, Drosophila melanogaster, and Saccharomyces cerevisiae, and simulated datasets. (Upper-right) Three examples of highly-skewed and zero-inflated distributions were chosen, characterized by long tails and a variance that increases with the mean. (Lower-right) Four examples of almost-symmetric distributions are shown. (**B**) A cartoon depiction of the likelihood model, which expects each genotype to be drawn from a different underlying Normal sampling distribution (grey) with a specific mean and variance that is dependent on a genotype’s genetic variants, the selection coefficients of those, and the additive or epistatic model combining those (C). Observations can be zero with a certain probability (zero inflation), and the offspring number is scaled to be relative to the mean of the overall distribution of genotypes in a population. (**C**) Cartoon depicting how the mean relative offspring number of a genotype would theoretically vary depending on the number of positive and negative mutations it carries under different additive or epistatic models of combining effects. Six different global epistatic models can be used to combine the effects of those mutations: base additive or multiplicative, and combinations of those with synergistic, diminishing returns, and no epistasis. Gray area indicates negative values, which have no meaning for fitness.

The choice of function for combining mutation effects is not merely a mathematical triviality but rather represents a hypothesis on how molecular processes affect the development of a trait and how these, in turn, are jointly affecting fitness (18). For instance, the choice of a multiplicative model of fitness made by many theoretical population geneticists can be justified by assuming that mutations decrease survivorship by a certain fraction, independently of each other. On the other hand, quantitative geneticists traditionally consider traits such as height, and thus may choose a model where mutations activate or repress independent growth pathways and may compensate one one another additively. The additive and multiplicative models are approximately identical when effect sizes (i.e. selection coefficients if the trait is fitness) are small, as the product terms of the effect sizes vanish, but can differ more substantially when there are mutations of moderate or large effects, particularly in the tails of the distribution.

Despite recent conceptual efforts to connect GWA results and population genetics and natural selection concepts (23), a common GWA framework to directly test different global additive, multiplicative, or epistatic models is still lacking (but see (24)). Here we developed NAPs (Non-Additive Polygenic models) for joint inference of effect sizes for *m* loci together with parameters modeling how the effects of loci combine (additive, multiplicative, etc.). These models are so-called global epistasis models because they do not parameterize interactions between individual loci directly, but rather model the interaction as a function of the total combined effect of all alleles in all loci (25) (**Fig. 1C**).

Our aim was to develop a flexible GWA-like model that could explain individual trait variation as a function of genomic variation using both additive and non-additive functions commonly used in evolutionary genetics. Much of the theory on epistasis models in population and quantitative genetics deals with fitness and natural selection, so we developed the NAP models to be especially suited for fitness traits (or proxies thereof). Fitness is a complex trait that is of special interest (natural selection is defined as genetic variation in fitness), but often is non-Normally distributed and, therefore, requires careful modeling (26). Here, we define the relative fitness of an organism as the expected total number of offspring produced in a lifetime divided by the population mean number of offspring. Relative fitness is then bounded between zero and *N*, with mean=1. The number of offspring of an individual can be thought of as arising from the convolution of a Bernoulli representing viability and a Poisson describing the offspring number, which is poorly approximated by a normal distribution even after various normalizations (26) (**Fig. 1A**). In particular, modeling of offspring number is often challenged by the inflation of zeros present in the dataset (**Fig. 1B**), which cannot be easily modeled by truncating the negative side of a normal distribution and allowing a point mass at zero equal to the CDF of the normal distribution at this point. We therefore model the normalized number of offspring from individual *y*_*j*_ with genotype *j* as being drawn from a mixture distribution: *y*_*j*_ ∼ (1 − *p*) N (*w*_*j*_ *w*_*j*_*b* + *a*) + *p δ*_0_ (**Fig. 1B**); where *w*_*j*_ represents the unobserved true fitness of a given genotype in a population, were different genotypes have different *w*_*j*_ fitness (e.g. distribution means in **Fig. 2B** vary for different genotypes); *a* and *b* are constants that allow either for homoscedastic Gaussian variance, when *b* = 0, or for variance to increase with the mean, *b* > 0 (e.g. distribution widths in **Fig. 1B** vary for different genotypes). Finally, independent of the genotype, the model allows for a fraction, *p*, of stochastic zeroes, often observed in real offspring distribution data and fitness assays (e.g. fraction at zero in **Fig. 1B**). This framework is flexible enough to accommodate a wide array of sampling distributions (**Fig. 1A**). Also, in the data sets we will consider, the number of offspring per individual is relatively large so the normal distribution of offspring per genotype should approximate the Poisson quite well. Each genotype, *j* = 1…*N*, is assumed to have a fitness *w*_*j*_ that is a function of the genotypic state in the genome-wide mutations affecting fitness. The functions we will use to describe the combined effects of mutations can be additive:

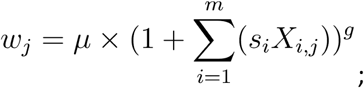

or multiplicative:

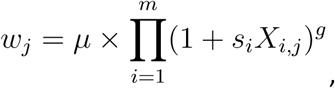

**Figure 2.**
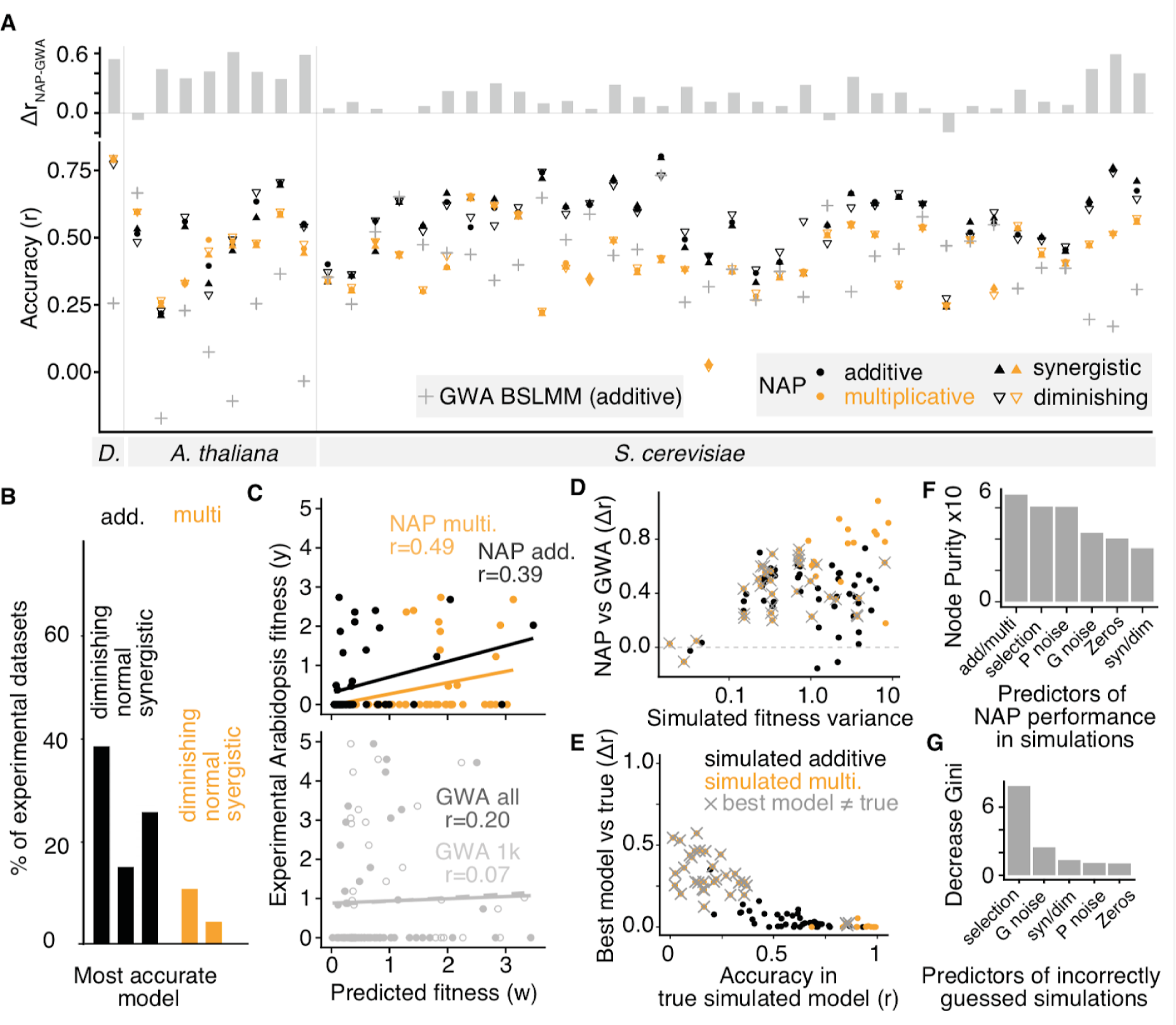
Significant non-additive selection in Arabidopsis, Drosophila and Saccharomyces experiments. **(A)** Cross-validation prediction accuracy using Non-Additive Polygenic (NAP) models in 44 fitness datasets in D. melanogaster (acronym D.), A. thaliana and S. cerevisiae. Accuracy was evaluated by Spearman’s r correlation coefficient between the actual and predicted value of relative fitness from the test set of individuals in each dataset. Fitness predictions were generated combining inferred selection coefficients for 1,000 SNPs using additive, multiplicative, or other non-additive functions. For reference, the prediction accuracy using state-of-the-art BSLMM Genome-Wide Association (GWA) (17) is also shown, along with the magnitude of accuracy improvement (Δr). **(B)** Percentage of the 44 datasets that were best predicted (highest r) by each of the six tested models. **(C)** Cross-validation predictions of relative offspring number in A. thaliana (experiment code “mli”, random test set n=52), with an example of the additive and the multiplicative models, and the comparison of predictability with BSLMM GWA model with all and the top 1,000 SNPs (random test set n=52). (**D-E**) Cross-validation prediction accuracy plots of 96 simulated datasets under a multiplicative (orange) or additive (black) polygenic model. Grey crosses indicate simulated multiplicative datasets for which the best model (highest r) was additive. (**D**) Each simulated dataset’s cross-validation accuracy improvement using NAP over BSLMM GWA, plotted against the fitness variance of each simulated dataset. (**E**) Each simulated dataset’s cross-validation accuracy difference between the true model (fitti ng NAP with the known parameters used to simulate the dataset), and the best model (NAP run with the parameters that maximized r). (**F**) A Random Forest was used to explain the accuracy difference between NAP and GWA based on the hyperparameters used to simulate datasets (n=96). Variable importance shows which simulation parameters lead to the largest changes in accuracy improvement. The parameters are: model type additive/multiplicative, selection strength, Poisson noise, Gaussian noise, random zeroes, synergistic/diminishing global epistasis (see text for explanation). (**G**) A Random Forest was used to explain which parameters (see F) characterize the simulated multiplicative datasets where the best model was additive (n=32).

Here *X* encodes the genotypes of biallelic SNPs, with alternative allele dosages: 0,1,2; assuming individuals are diploid, and *μ* is the fitness of a hypothetical reference genotype (with *X*=0 in all loci). Both additive and multiplicative functions can have different functional forms that make them depart from linear or geometric functions when *g* ≠1. Specifically, when *g* > 1, the genotype-trait architecture is said to have synergistic epistasis. When *g* < 1, it has diminishing returns epistasis. In total, six combinations of model architectures can be tested: purely additive, additive synergistic, additive with diminishing returns, purely multiplicative, multiplicative synergistic, and multiplicative with diminishing returns (**Fig. 1C**). Given the above sampling distribution per genotype and parameters, the likelihood of observing relative offspring number *y* of a genotype *j* is:

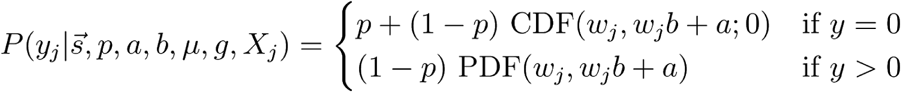

And the log-likelihood of all observations is the sum over all log-likelihoods per genotype:

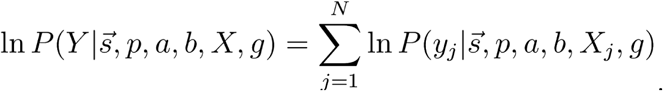

The key predictor in our NAP model is the vector of relative fitness deviations or selection coefficients on genetic variants, *s* ∈ [−1, *Inf*]. Following (27), we define a selection coefficient as the relative fitness advantage or disadvantage of a minor allele with respect to the fitness of a reference haplotype (the theoretical haplotype with all major alleles).

To infer the vector *s* and the model hyperparameters, we use the multivariate quasi-Newtonian optimization algorithm Spectral Projected Gradient (SPG) (28). Although this procedure has been developed for large-scale multivariate optimization and is implemented in C++ (https://github.com/MoisesExpositoAlonso/napspg), conducting these analyses with millions of SNPs is computationally-demanding and time-consuming. Hence, we instead first run a Bayesian Sparse Linear Mixed Model (BSLMM, implemented in GEMMA) (12), a polygenic additive GWA model for which efficient optimizations have been implemented. Using BSLMM, we can pre-select a set of top associated SNPs with the trait for which we then run our likelihood optimization. The SPG optimization was run until convergence or for 2,500 iterations. To assess goodness-of-fit and avoid overfitti ng, we use a cross-validation approach, where 90% of the data is used for training and 10% is used for testing. Accuracy is measured using Spearman’s rank correlation (r) between predicted and measured values of relative offspring number of individuals.

Using the NAP model, we analyzed real fitness datasets and tested which polygenic architecture model best fits the data. We used datasets from three model organisms: *Arabidopsis thaliana* (part of the 1001 Genomes Project (2)), *Drosophila melanogaster* (part of the Drosophila Genetic Reference Panel (DGRP2) (3, 29)), and *Saccharomyces cerevisiae* (part of a >1000 genomes effort (30)). For *A. thaliana*, we analyze a total of eight experimental datasets, 515 natural lines, and 4,438,427 biallelic SNPs. For *D. melanogaster*, we use one dataset, 205 natural lines, and 4.5 million biallelic SNPs. For *S. cerevisiae*, we use 35 datasets, 1011 natural lines, and 83,794 biallelic SNPs. BSLMM Genome-Wide Associations for the three species confirmed the results from the original publication that fitness traits were polygenic, with the highest estimated number of causal loci (*nγ*) for any of the analyzed environments (mode of 1,000 MCMC steps) being: 63 for *Drosophila*, 261 for *Arabidopsis*, and 271 for *Saccharomyces.* None of the previous publications tested whether these polygenic traits were additive or not. For each of these 44 datasets we applied the likelihood method to the 1,000 SNPs with highest posterior probability in BSLMM, for different (non-)additive architectures, and selected the best models based on cross-validation accuracy (**Fig. 2**).

Our results show that non-additive polygenic (NAP) models in general tend to provide higher prediction accuracy than standard additive models (**Fig. 2A-B**). The most common architectures were either additive with diminishing global epistasis or additive with synergistic global epistasis, and a significant 15% fraction were also best fit under a multiplicative model in at least one environment for each of the three species.

As expected due to the zero-inflation and long tails in these datasets (**Fig. 1A**), the state-of-the-art Genome-Wide Associations had a lower prediction accuracy than the NAP models (**Fig. 2A-C**) for almost all data sets. The NAP model improved cross-validation accuracy over GWA by an average of r=0.216 (2.5-97.5% quantiles: 0.0729, 0.592). This was more pronounced when the best model was multiplicative (Δr=0.32 multiplicative, Δr=0.35 multiplicative with diminishing returns) compared to additive (Δr=0.21 for additive, Δr=0.16 for additive diminishing, Δr=0.23 for additive synergistic). To exclude that these improvements were due to the winner’s curse, we re-ran the GWA with the top 1,000 SNPs from the first GWA run which were used to fit NAP (for comparison of the all SNPs GWA and the top 1,000 SNPs GWA, see **Fig. 2C**).

To test the robustness of our conclusions, we ran the same set of analyses in 96 simulated datasets (**Fig. 1A**). The fitness of 1,500 individuals was simulated using 10,000 biallelic SNPs in 500 causal loci, each randomly assigned a selection coefficient assuming *log*(1 − *s*) ∼ 𝒩 (0, *σ*); with variance either 0.01 (weak selection) or 0.1 (stronger selection). The strength of selection can also be interpreted as the polygenicity of the trait, as in the weak selection case only ∼30% of SNPs have selection coefficients > 1%, while in the strong selection case >90% of the 500 loci have selection coefficients > 1%. Individual-level fitness is then calculated as a combination of all genome-wide selection coefficients using an additive or multiplicative formula (see above), and in all combinations with or without synergistic and diminishing returns epistatic parameters (using a grid of parameter values from 0.8 to 1.2 in 0.1 increments). Finally, we added different degrees and types of sampling noise, either 0% or 20% zero-inflation of random zeroes, a basal Gaussian noise of 0.01 or 0.5 (*a*), and variance increased in proportion to the mean of 0.01 or 0.5 (*b*); and all combinations of the above (n=96), creating an array of fitness distribution shapes. The hyperparameter values were manually selected to create realistic fitness distributions similar to those observed in *A. thaliana, D. melanogaster* and *S. cerevisiae* datasets (**Fig. 1A)**.

Using the same approach as with the experimental data, we conducted a pre-selection of the top SNPs using BSLMM, ran our NAP models with 1,000 SNPs under different additive and multiplicative architectures and a grid of values of the epistatic parameter, and then selected the best models based on cross-validation accuracy. As before, we found that our best model had an average cross-validation predictability of r=0.50, while GWA had an average accuracy of r=0.21. The improvement in accuracy of NAP over GWA increased as the fitness distribution became more complex, that is, heavy-tailed (skewness, Pearson’s r=0.41, *P=* 3.64×10^−5^) and broad (variance, r=0.24, *P=* 1.89×10^−2^) (**Fig. 2D**).

Because the true underlying genome architecture is known for the simulated datasets, we could study the performance of our likelihood optimization and cross-validation approaches, and compare them with the BSLMM GWA baseline. We found that 98% of all additive simulated datasets had the highest prediction accuracy under an additive model. Fitting different *g* global epistatic parameters (0.8-1.2 in 0.05 increments) in these additive models and selecting the model with highest prediction accuracy, we nevertheless identified the correct simulated epistatic parameter 20% of the time, close to the random expectation 10%. This may be caused by the minor influence of this global epistasis parameter in shaping fitness (see **Fig. 2 F**), or due to optimization issues and/or our coarse grid search approach.

Multiplicative architectures did not tend to improve prediction accuracy even when the true model was in fact multiplicative. Only 32% of multiplicative simulations found highest prediction accuracy under a multiplicative model. When we studied which multiplicative datasets had poor prediction accuracy under a multiplicative model, we found that it was those datasets with high noise, leading to an overall low accuracy of the model (**Fig. 2D-E**). We conducted a Random Forest analysis to quantify the simulation conditions that characterized runs where the wrong model had the highest prediction accuracy. Unsurprisingly, it was simulations under conditions with weak multiplicative selection and high Gaussian noise that were wrongly identified as additive (**Fig. 2F-E**, crosses), as in cases where loci have small effects, additive and multiplicative models are approximately identical. Despite the fact that simulated multiplicative datasets were consistently better predicted with additive rather than multiplicative models, we found that additive models fitted with synergistic global epistasis parameter (*g*≥1.2) had on average a 6% higher accuracy than additive models with diminishing returns parameter (*g*≤0.85) (t-test, t = 2.433, df = 189.7, p-value = 0.0158; Wilcoxon rank sum test, W = 5631, *P* = 0.0079). This shows that our algorithm may detect non-linear multiplicative-like architectures, but they appear to be well approximated by an additive model with synergisms (see the similarity in multiplicative and additive-synergistic lines in **Fig. 1C**).

Here we presented the NAP model, the first Genome-Wide Association pipeline that enables researchers to choose a global non-additive architecture in their association study for prediction purposes. Applying NAP to three diverse species, we find evidence for a multiplicative genotype-fitness architecture at least in some environments, and simulations suggest our model selection based on prediction accuracy is conservatively biased towards additive architectures. Beyond testing different polygenic architectures, the tailored likelihood model behind NAP accommodates non-Normal phenotypic distributions and achieves a higher cross-validation accuracy when predicting trait values of genotypes hidden from the model, which could have broad applications both to study natural selection and fitness, as well as other traits. We also note that the method has potential for improving phenotypic prediction in human disease studies and in other studies that rely on the standard additive polygenic risk score model (12).

Conclusions on global epistasis could arise due to non-linearities in the measuring scale of a trait (31). Fitness has a natural scale, i.e. the number of offspring produced by an individual, and therefore should not be affected by this confounding. However, some proxies for fitness, e.g. growth rates for unicellular organisms, could also be problematic for measuring selection (32).

The evidence for non-additive genetic architecture presented here may not be surprising given the growing literature of experiments that require genome-wide epistasis to explain asymmetric responses to artificial selection (33), line-dependent effects of mutations in *Drosophila* (14, 34), or significant quantitative trait loci hubs in yeast (35). This has sparked recent development of various statistical approaches to test epistasis more generally, by studying the emergent patterns of epistasis as its contribution to variance (36), or one genotype-to-trait map (24, 25, 31, 37, 38) as in this study. Recent systems biology approaches for creating massively-parallel mutations using CRISPR/Cas9 techniques (39), as recently aimed in yeast experiments (40), should further enable researchers to test the underlying additive or epistatic interactions of mutations. The evidence for non-additive genetic architecture in three key model species identified in this study will have substantial impact in our understanding and predictability of polygenic adaptation of species (6, 41, 42).

## Acknowledgements

We are thankful to Graham Coop for pointing us to relevant references, to Dmitri Petrov, Ben Good, Patricia Lang, Clemens Weiss, and all the members of the Moi Lab for discussions and/or comments on the manuscript. This work was supported by EMBO and Carnegie Endowment Funds (M.E.-A.) and the National Institutes of Health (R01GM116044; R.N.). Some of the computing for this project was performed on the Memex and Calc clusters. We would like to thank Carnegie Institution for Science and the Carnegie Sci-Comp Committee for providing computational resources and support that contributed to these research results.

## ADDITIONAL INFORMATION

### Author contribution

M.E.A and R.N. conceived the project. M.E.A, P.W., and R.N. developed the methods. M.E.A. conducted the analyses, and wrote the software. M.E.A, P.W., and R.N. wrote the manuscript.

### Data and code availability

All code and method implementation details are available at https://github.com/MoisesExpositoAlonso/napspg with doi: 10.5281/zenodo.3934749.

### Disclosure statement

M.E.A and R.N. declare no competing financial interests. P.W. is currently employed by 23andMe. The funders of this study had no role in the study’s design, data collection and analysis, decision to publish, or preparation of the manuscript.

